# BrAIn: A comprehensive artificial intelligence-based morphology analysis system for brain organoids and neuroscience

**DOI:** 10.1101/2025.02.19.638973

**Authors:** Burak Kahveci, Elifsu Polatli, Ali Eren Evranos, Hüseyin Güner, Yalin Bastanlar, Gökhan Karakülah, Sinan Güven

## Abstract

Human-induced pluripotent stem cells (iPSCs) offer transformative potential for biomedical research, with iPSC-derived organoids providing more physiologically relevant models than traditional 2D cell cultures. Among these, brain organoids are particularly valuable for drug screening, disease modeling, and investigations into molecular pathways. Accurate representation of brain morphology is critical, as more complex organoid structures better mimic the human brain. Deep learning (DL) and machine learning (ML) approaches have become integral to analyzing organoid morphology, yet tools for comprehensive, time-resolved assessments are scarce. Here, we introduce **BrAIn**, a DL-based application for analyzing the developmental progression of brain organoids. BrAIn tracks their evolution from embryoid bodies and quantifies parameters including area, Feret diameter, perimeter, roundness, and circularity. It also classifies budding and abnormal morphologies of 3D organoids and detects monolayer neural rosette structures, key features of neuronal differentiation. Designed with accessibility in mind, BrAIn provides a no-code interface, enabling researchers of all technical backgrounds to conduct advanced morphological analyses with ease. Our study demonstrates the application of BrAIn to evaluate the effects of different growth conditions - static, orbital shaker, and microfluidic chip-based - on brain organoid development. Orbital shaker cultures resulted in the largest organoids, while chip-based systems achieved more homogeneous growth. Both conditions produced organoids with greater morphological complexity compared to static culture. BrAIn emerges as a robust, user-friendly tool to quantify brain organoid development and explore how versatile growth conditions influence their morphology and maturation.

## 1. Introduction

Human-induced pluripotent stem cells (iPSCs) have become one of the most promising tools for medical research offering immense capacity for obtaining cellular phenotypes [1]. iPSCs can also give rise of organoids, self-formed 3D tissue mimicries, providing the structural and functional recapitulation of an organ [2]. iPSCs derived-brain organoid can be used in modeling neurodegenerative diseases [3, 4], drug discovery research [5], or platforms [6–9] for determination of neurotoxicity [10]. Morphology of brain organoids can be potential indicator of physiologically and functionally relevant tissue mimicry. Besides numerous advantages provided by brain organoids, time and cost spent can be significant burdens even for well-established research groups and enterprises [11]. To improve brain organoid generation process, increase the yield, and minimize human-based errors, experimental planning, optimization of resources, and interpretation of results should be well defined [12].

Machine learning models are used in many areas such as decision support systems in medicine, analysis of medical images, disease diagnosis and prognosis prediction [13]. Deep learning methods provide automatic feature extraction, and they have the advantage of exploiting GPUs for parallel operations on diverse data such as image, text, sound and genomic [14, 15]. Image datasets pioneered deep learning methods providing diverse scenarios such as image classification, segmentation, object detection and tracking in microscope images [16].

Abundant and heterogeneous data assist in generation and reflection more realistic, real-life scenarios. Images from wet lab experiments are laborious, costly and limited in number, restraining capacity for deep learning models. Introducing data augmentation methods, where artificially modified inputs are generated, promise to overcome this problem providing unlimited data. Traditional data augmentation methods include operations such as flipping, cropping, rotation, and color space transformations [17]. However, traditional data augmentation methods are often limited in improving model performance [18, 19]. To overcome the limitations of traditional data augmentation methods, generative adversarial networks [20] are used for data augmentation [21–23]. Generative adversarial networks create new images based on the given datasets [24]. StyleGAN3, an approach based on generative adversarial networks, is used in the medical field and life sciences [25, 26].

Deep learning applications in microscope images obtained from organoids can both facilitate the examination of images and provide the opportunity to optimize the experimental processes against human errors[27]. In addition to classification, object detection and image segmentation are extensively applied in organoid analysis performed with deep learning [27]. Transfer learning is a deep learning approach that leverages pre-trained models to achieve better accuracy with a limited amount of data [28]. This approach has proven to be highly effective to improve the performance measured by classification, detection, and segmentation evaluation metrics [29, 30].

Deep learning applications offer quite promising opportunities in the field of medical computer vision [31]. Morphological studies on stem cells and iPSCs have been conducted using deep learning [32–36]. Zhu et al. used convolutional neural networks to characterize neural stem cell differentiation using brightfield microscope images [33]. Kahveci et al. used YOLOv5, a deep learning-based object detection model, to detect spermatogonial stem cells from mouse testes with a complex microenvironment [34]. The DenseNet121 model was used by Kim et al. to determine the differentiability of human adult stem cells obtained from different donors [36]. Also, there are many image processing and deep learning-based studies for use in morphological analysis of organoids [37]. These studies can be divided into three main areas: classification [38–42], segmentation [43–49], and detection [50–52].

There exist several previous studies on organoid classification with deep learning. Abdul et al. developed a deep learning model that classifies colon organoid images in terms of morphological features such as opacity and budding [38]. To automatically evaluate drug screening of organoids, Bian et al. developed a system combining CNN-based feature extraction, multi-head classifier, and fully connected network [39]. Park et al. developed a system to predict kidney organoid differentiation status using DenseNet121 [40]. Huang et al. used a deep learning-based model to identify cystic and solid morphologies in colorectal cancer organoids [41]. Kegeles et al. developed a transfer learning-based model to analyze the differentiation of retinal organoids [42]. Asano et al. developed a model that predicts whether differentiation is progressing appropriately in hypothalamic-pituitary organoids using a deep learning approach [53].

Deep learning and image processing methods are employed in morphological analyses of organoids frequently. Borten et al. used a combination of smoothing and adaptive thresholding methods for morphological analyses of spheroids and organoids [43]. The created application was used for morphological analyses of breast-cancer spheroid, colon, and colorectal-cancer organoid images. Gritti et al. developed a deep learning-based model for morphological and fluorescence analyses of pescoids and brain organoids [44]. Matthews et al. used U-Net to analyze parameters such as count and size of organoids [45]. They trained this model on pancreatic cancer organoid images and tested it on pancreatic, lung, colon, and adenoid cystic carcinoma organoid images. Park et al. used U-Net to perform morphological analyses of colon organoids [46]. They measured many parameters such as area, perimeter, and diameter with the developed model. Lefferts et al. trained a Mask-RCNN-based model for the analysis of some morphologies in patient-derived organoids [47]. Swelling measurements were made in the organoids with this model. Wang et al. developed a deep learning-based model that segments retinal, cardiac, and brain organoids and their internal structures using optical coherence tomography images [48]. They measured parameters such as size, area, and volume with this model. Chen et al. trained a modified U-Net model for tumor spheroid analysis [49]. They performed tumor boundary detection and invasiveness analysis with this model. Deben et al. developed a deep learning-based analysis tool for lung and pancreas drug screening in patient-derived organoids [54]. This tool can perform area measurement. Deben et al. also developed a metric called Normalized Organoid Growth Rate (NOGR) metric for pancreatic organoid analysis and used it for segmented images obtained with a deep learning-based model [55].

Bian et al. used deep neural networks for organoid tracking and detection [50]. The model was trained on z-stack images obtained from a brightfield microscope. Leng et al. used a YOLOX-based model to detect intestinal organoids [51]. The model was trained on brightfield microscope images. Kassis et al. used a Faster R-CNN model for intestinal organoid detection and tracking [52]. This model was trained on brightfield microscope images and was able to measure the size distribution of organoids.

There are a few studies on the classification and morphological analysis of neural organoids. For the prediction of neurotoxicity and the state detection of Parkinson’s disease using the midbrain organoid, Monzel et al. performed a phenotypic-based analysis study using the Random Forest model [56]. With this model, toxicity was predicted with 86% accuracy and disease state was predicted with 93% accuracy at day 70. Albanese et al. developed an analysis system based on image processing and deep learning with cerebral organoid datasets obtained from fluorescent microscopy [57]. This system performed both single cell and cytoarchitecture analyzes of cerebral organoids. The system has also been used on Zika virus-infected organoids. Metzger et al. used a deep learning model for drug screening on neural organoids [58]. Applying the method to Huntington’s disease, they showed that bromodomain inhibitors can reverse the complex phenotypes associated with the disease and that this target may be a potential avenue for drug development. Deininger et al. used brain organoid MRI images to study organoid growth with the U-Net model and identified cystic and noncystic organoids with high accuracy [59]. Brémond-Martin et al. used U-Net, K-means, SVM methods to characterize the morphological development of brain organoids. They identified morphological patterns specific to day 14 neuroepithelial formations [60]. Brémond-Martin et al. proposed MU-Net, a scaled-down version of U-Net architecture, for the segmentation of brain organoids (BOs) with small datasets and compared it with different data augmentation strategies. MU-Net exhibited a more robust performance compared to U-Net, achieving segmentation results with almost the same accuracy as optimized augmentations on small datasets [61].

Neural organoid studies using machine learning and deep learning models are presented in Table 1.

**Table 1.** Classification and segmentation studies using neural organoids.

|  | **Dataset** | **Method** | **References** |
| --- | --- | --- | --- |
| **Classification** | Midbrain Organoids | Random Forest | [56] |
|  | Neural Organoids | Convolutional neural networks | [58] |
| **Segmentation** | Cerebral Organoids | U-Net | [57], [59] |
|  | Brain Organoids | U-Net, K-means, SVM | [60] |
|  | Brain Organoids | Mu-Net | [61] |

Organoid morphology analysis and developmental morphology monitoring are quite critical. Although semi-automated microscope applications provide a certain level of analysis, they are still time-consuming. In addition, since human intervention is required, there may be potential incorrect measurements and errors. As summarized in Table 1, there are studies that perform organoid morphology analysis. However, there is no comprehensive tool that can analyze the entire developmental process. In this study, the BrAIn tool was developed to analyze the developmental process of brain organoids. Morphological analyses of the developmental process of the brain organoid can be analyzed and quantified with BrAIn under three different sections.

- In the classification section, three different classes of morphologies (normal, abnormal, and budding) that may occur during the process of creating brain organoids from embryoid bodies can be classified with BrAIn.
- BrAIn can segment embryoid bodies and brain organoids with high precision, and the developmental processes of brain organoids can be quantified with different parameters. Developmental processes were examined under different growth conditions. The different effects of these conditions on growth and morphology were analyzed with BrAIn.
- In addition, neural rosette structures were identified, which are quite important in the process of generating iPSC-derived neurons and are difficult to detect with the naked eye.

With BrAIn, every step of the brain organoid generation process is automated, minimizing human-based errors, saving time and decreasing cost by planning the experimental processes based on artificial intelligence, all through a user-friendly interface that even users with no coding knowledge can easily navigate.

## 2. Methods

### 2.1. hiPSCs Culture

Human induced pluripotent stem cells (hiPSCs) are kindly provided by Prof. Tamer Onder [62]. hiPSCs were cultured in 6-well plates coated with 1% embryonic stem cell (ESC) qualified Matrigel (Corning) with mTeSR1 (STEMCELL Technologies) medium. Culture was maintained with a daily change of fresh medium until cells reached 80% confluency. hiPSCs were passaged with ReLeSR (STEMCELL Technologies) at a ratio of 1:10.

### 2.2. Neuronal Differentiation

hiPSCs were differentiated into NPCs according to previously published protocol [63] using dual SMAD inhibition. Briefly, iPSCs detached with ReleSR and seeded on low attachment 6-well plates in human embryonic stem cell medium (hESCM) (DMEM/F-12 (Gibco, Thermo Fisher Scientific), 20% knockout serum replacement (KOSR) (Gibco, Thermo Fisher Scientific), 1% non-essential amino acids (NEAA) (Lonza), 1% non-essential amino acids (NEAA) (Lonza), 1% GlutaMAX supplement (Gibco, Thermo Fisher Scientific), 1% penicillin/streptomycin, 100 mM 2-mercaptoethanol with the fresh addition of ROCKi (50 µM) for 2 days. After 2 days, medium was replaced with SMAD inhibitors using with 5 µM SB 431542 and 10nM LDN 193189. Embryoid bodies were incubated for further 2 days. At day 4, medium was replaced with Neural induction medium (NIM) (DMEM/F-12, 1% N2, 1% NEAA, 1% penicillin/streptomycin, and 2 µg/ml heparin). Embryoid bodies were incubated as suspension for 2 days, then seeded on growth factor-reduced Matrigel (Corning) coated wells with the NIM for the rosette formation. Media were changed every 2 days until neural rosette structures appeared.

### 2.3. Development of Brain Organoids

hiPSCs were differentiated into brain organoids according to previously published protocol [6]. Briefly, hiPSCs were removed from Matrigel with ReleSR and dissociated with pipetting gently. Cells were seeded on low attachment U-bottom 96-well plates at a density of 9×10^3^ cells/well with embryonic stem cell medium (hESCM) with the addition of ROCKi (50 µM) and 4ng/ml bFGF.

Cells were incubated for 6 days with half of media change every other day. Embryoid bodies were transferred into low attachment 24-well plates with Brain organoid Neural Induction Media (BONIM) (DMEM/F-12, 0.01% N2, 1% Glutamax,1% penicillin/streptomycin, and 2 µg/ml heparin) for 4 days. EBs were embedded into basement Matrigel (Corning) with Cerebral Organoid Differentiation Media (CODM) without the addition of vitamin A (1:1 DMEM/F12 and Neurobasal Medium (Gibco, Thermo Fisher Scientific, 0.5X N2, 1X GlutaMAX, 0.5X NEAA, 1X B27 w/o VitA, 0.25 (v/v) insulin, %0.02 (v/v) 2-mercaptoethanol). After 4 days, immature organoids were transferred into microfluidic chips, orbital shaker, and static conditions according to their groups with the addition of vitamin A into CODM. Cells were incubated for 48 days with medium changes 2-3 times a week (Figure 1A).

**Figure 1.**
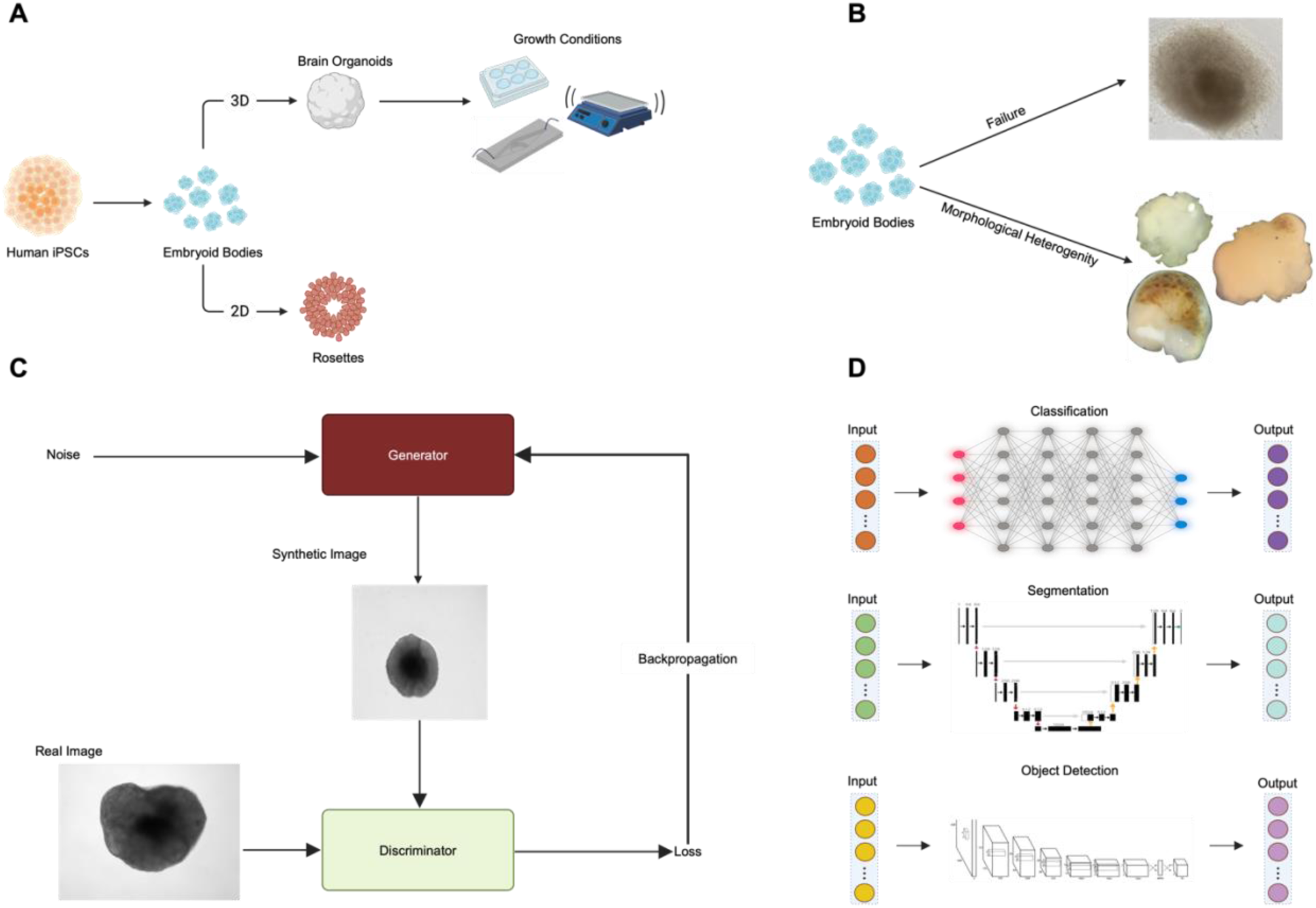
Workflow summarizing experimental and computational stages of BrAIn. A) Schematic illustration showing generation of Brain Organoids and Rosettes derived from hiPSCs. B) Brain organoids originate from the same embryoid body stage but have high heterogeneity. C) Schematic diagram of the StyleGAN3 model used in data augmentation to ensure better results and generalization of BrAIn. D) Illustration of the deep learning models used for the three sections included in BrAIn, classification, segmentation and object detection.

### 2.4. Fabrication of Microfluidic Chips

Microfluidic chips consist of multilayer poly (methyl methacrylate) (PMMA). Channels were designed by AutoCAD (Autodesk) and CorelDRAW (Corel Co.) (Figure S1). PMMA layers were cut with a laser cutter (Epilog-MINI) and attached by double-sided adhesive. Sterilization of microfluidic chips was performed with 70% ethanol following UV exposure for 30 minutes. Brain organoids were cultured in the microfluidic chip with 35 µl/min.

### 2.5. Immunofluorescence Staining

Embryoid bodies and brain organoids were fixed with 4% paraformaldehyde (Sigma-Aldrich) for 24 hours at 4 °C. Brain organoids were embedded in cryomatrix (OCT, Fisher Healthcare) and cryosections were taken with Leica CM1950 (Leica Biosystems, Germany). Embryoid bodies and brain organoids were permeabilized with permeabilization buffer (0.3% Triton X-100 (neoFroxx, GmbH, Cat no. 8500) (v/v) in PBS (Gibco, Thermo Fisher Scientific, Cat no. P04-36500) for 15 min at room temperature. Cells were treated with blocking buffer (0.05% Tween 20 (Sigma-Aldrich, Cat no. P7949) (v/v), 1% Bovine Serum Albumin (BSA; Sigma-Aldrich, Cat no. A2153) (w/v) in PBS) for 15 min at room temperature. Primary antibodies (α-SMA (Cell Signaling Technology, Cat no. 48938S), Nestin (Proteintech, Cat no. 19483-1-AP), Sox17 (Abcam, Cat no. Ab84990), Sox2 (Bioss, Cat no. bs-0523R), Tuj1 (R&D Systems, Cat no. MAB1195), and N-cadherin (Cell Signaling Technology, Cat no. 13116T) were added to the relevant samples and incubated overnight at 4 °C. Cells were stained with relevant secondary antibodies for 2 h at room temperature. Nuclei were stained with 4,6-diamidino-2-phenylindole (0.5 μg/mL) (DAPI; Neofroxx, Cat no. 1322). Images of the samples were captured with a fluorescence microscope (Olympus IX71) and a confocal microscope (Zeiss LSM880).

### 2.6. Datasets and Training of Models

In the training process of BrAIn, five datasets were used: two datasets containing normal-abnormal classes and normal-budding classes for the classification section (Figure 1B), time-dependent brain organoid and embryoid body datasets for the segmentation section, and a rosette dataset for the object detection section.

Details of the dataset and model information are given in Table 2. In addition, the brain organoids dataset in the segmentation section and the rosette datasets in the object detection section were augmented using the StyleGAN3 model [64]. The working process of the Generative Adversarial Network on which StyleGAN3 is based is given in Figure 1C. As summarized in Figure 1D, BrAIn contains three different parts: classification, segmentation, and object detection. Transfer learning approach was used in classification. In this section, 5 models were used: DenseNet [65], Inception [66], ResNet50 [67], VGG16 [68] and Xception [69]. In the segmentation part, the U-Net [70] model was used for the segmentation of brain organoids and embryoid bodies. In the object detection part, the YOLOv8 [71] model was used to detect rosette structures from 2D cell culture images.

**Table 2.** Dataset and Model information for BrAIn.

| Model | Dataset Name | Section | Train & Test Data Distribution | Epoch | Batch Size | Loss |
| --- | --- | --- | --- | --- | --- | --- |
| ResNet50 | Budding-Normal | Classification | 265-67 | 50 | 16 | Categorical Cross Entropy |
| DenseNet121 | Abnormal-Normal | Classification | 188-52 | 50 | 32 | Categorical Cross Entropy |
| U-Net | Embryoid Body | Segmentation | 132-33 | 20 | 4 | Binary Cross-entropy |
| U-Net | Brain Organoid | Segmentation | 1273-318 (898 Synthetic Images) | 10 | 4 | Binary Cross-entropy |
| YOLOv8 | Neural Rosette | Object Detection | 338-35 (293 Synthetic Images) | 100 | 8 | Distribution Focal Loss |

In classification model training step, the batch size was determined as 32 and the resolution of the images was 550×550. Early stopping approach was used throughout the training. Training was terminated if no improvement was observed in the validation loss value for three epochs. In the segmentation part, U-Net was used for the segmentation of brain organoids and embryoid bodies. Datasets for U-net were prepared with OrganoLabeler developed by our group [72]. In the models, the batch size was determined as 4 and the learning rate value was used as 1e-3. Cross validation approach was applied during the training process of the models and the fold value was determined as 5. Two brain organoids developed from static, microfluidic chip and orbital shaker systems were used for morphology analysis. In the object detection section, the YOLOv8 model was used to detect neural rosettes. In model training, the batch size was determined as 8 and the initial learning rate was used as 0.01. The training processes and parameter details of the models are given in Table 2. All datasets used were captured from an inverted microscope.

## 3. Results

### 3.1. Characterization of neural rosettes and brain organoids

Brain organoids have previously been established under dynamic conditions, such as orbital shaker and microfluidic chip systems, alongside traditional static growth methods. To comprehensively evaluate the performance of BrAIn across diverse culture conditions, brain organoids were generated in three distinct environments: microfluidic chip, orbital shaker, and static culture. Phenotypic characterization of brain organoids was performed with immunofluorescence staining, validating their differentiation potential through demonstrating presence of neural progenitor (SOX2) and neuronal cells (TUJ1) (Figure 2A-C).

**Figure 2.**
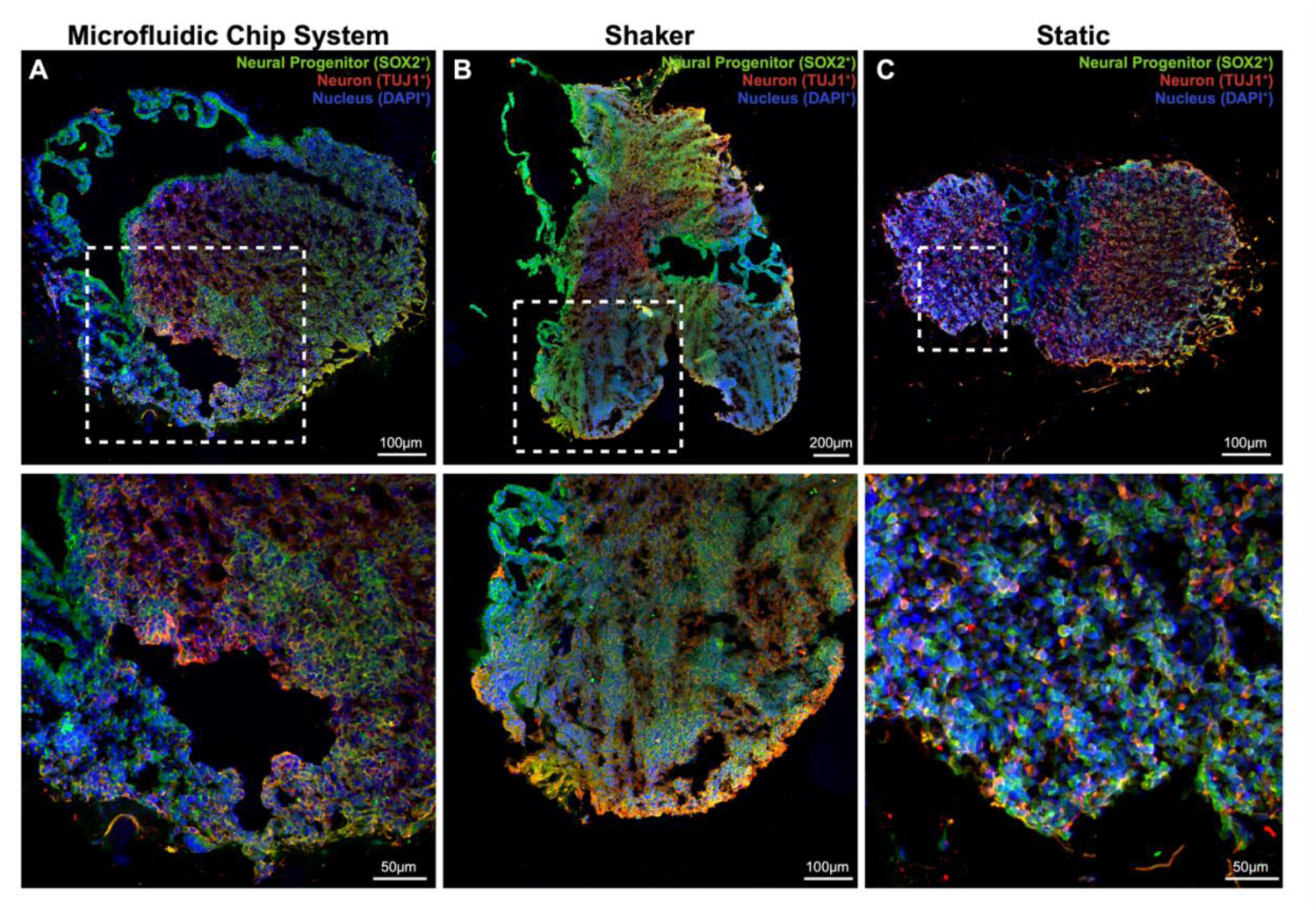
Immunofluorescence characterization of Day 30 brain organoids demonstrating SOX2 (green) expressing neural progenitor cells and TUJ1 (red) positive neurons. Cell nuclei were stained with DAPI (blue). A) Brain organoids grown in a microfluidic chip system. B) Organoids cultured in an orbital shaker. C) Brain organoids under static conditions. The lower panel provides a higher magnification view of the highlighted area.

Embryoid bodies (EBs) mimic early embryogenesis and have the potential to differentiate into the three germ layers, notably the ectoderm layer, which gives rise to the brain tissue. In this study, we generated EBs from hiPSCs using a 6-day suspension culture and confirmed their trilineage differentiation potential into ectoderm, endoderm, and mesoderm (Figure S2). Neural rosettes, formed during ectodermal differentiation, contain neural progenitor cells (NPCs) capable of differentiating into neurons, oligodendrocytes, and astrocytes—the primary neuroectodermal cell types. To enable BrAIn to accurately detect neural rosettes, which are challenging to identify through visual inspection, we developed hiPSC-derived neural rosettes and validated their structural integrity and differentiation potential using immunofluorescence staining (Figure S2).

### 3.2. Classification of brain organoid morphologies

Morphology plays a crucial role in determining the quality of brain organoids during their transition from embryoid bodies to organoids. Perturbation of organoid morphology at early-stage, formation of budding structures, serve as key indicators of successful brain organoid differentiation.

The classification module of BrAIn categorizes organoids into three distinct classes: normal, which lacks budding yet is considered acceptable; budding, characterized by the presence of perturbated zones; and abnormal, where cells fail to establish the basic structural integrity required for proper embryoid body formation (Figure 3A). BrAIn focuses on two primary classification tasks: distinguishing normal from abnormal organoids and differentiating between normal and budding organoids. For both tasks, transfer learning models pre-trained on the ImageNet dataset were utilized, employing five different deep learning architectures. The dataset was split into 80% training and 20% testing, and a 5-fold cross-validation approach was applied to ensure robustness.

**Figure 3.**
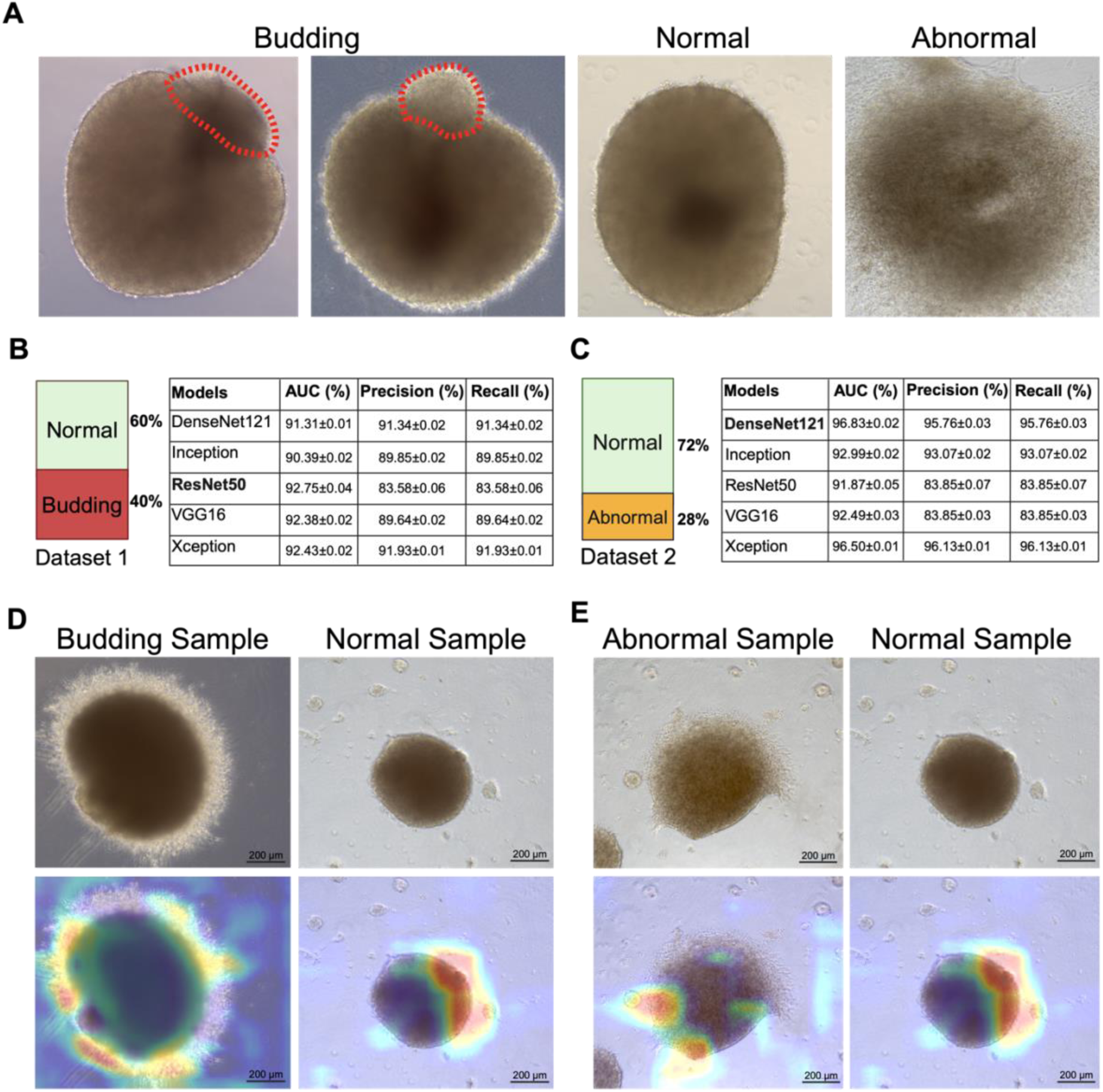
BrAIn’s classification module for brain organoids. A) Morphological features used for classification tasks, including Normal, Budding, and Abnormal categories. B) Class distribution and model performance metrics for Normal vs. Budding classification. C) Class distribution and model performance metrics for Normal vs. Abnormal classification. D) Grad-CAM heatmap illustrating the feature distribution utilized by the model for Normal vs. Budding classification. E) Grad-CAM heatmap illustrating the feature distribution utilized by the model for Normal vs. Abnormal classification. The red regions indicate high importance, highlighting the key areas used by the model during the classification process.

In the normal-budding classification task, the dataset consisted of 60% normal and 40% budding samples. Among the tested models, ResNet50 achieved the highest performance, with AUC, precision, and recall values of 92.75%, 83.58%, and 83.58%, respectively (Figure 3B). For the normal-abnormal classification task, which had a class distribution of 72% normal and 28% abnormal samples, DenseNet121 outperformed other models with an AUC of 96.83%, precision of 95.76%, and recall of 95.76% (Figure 3C). Grad-CAM heatmaps were employed to visualize the regions most influential in the model’s decision-making process, with red areas indicating the most significant features used for classification (Figure 3D, 3E) [73].

### 3.3. Brain organoid segmentation and morphological evaluation

To accurately capture the three-dimensional structure of brain organoids, the z-stack imaging method was employed during dataset creation, with a layer spacing of 10 µm (Figure 4A). OrganoLabeler, a labeling tool developed by our group, was utilized to annotate both the embryoid body and brain organoid datasets (Figure 4B, 4C). Throughout the embryoid body formation process, a segmentation model was developed based on images collected over four days, allowing for a detailed analysis of morphological changes. To further assess brain organoid development, images were captured at 7-day intervals from Day 20 to Day 62, providing comprehensive insights into organoid growth dynamics.

**Figure 4.**
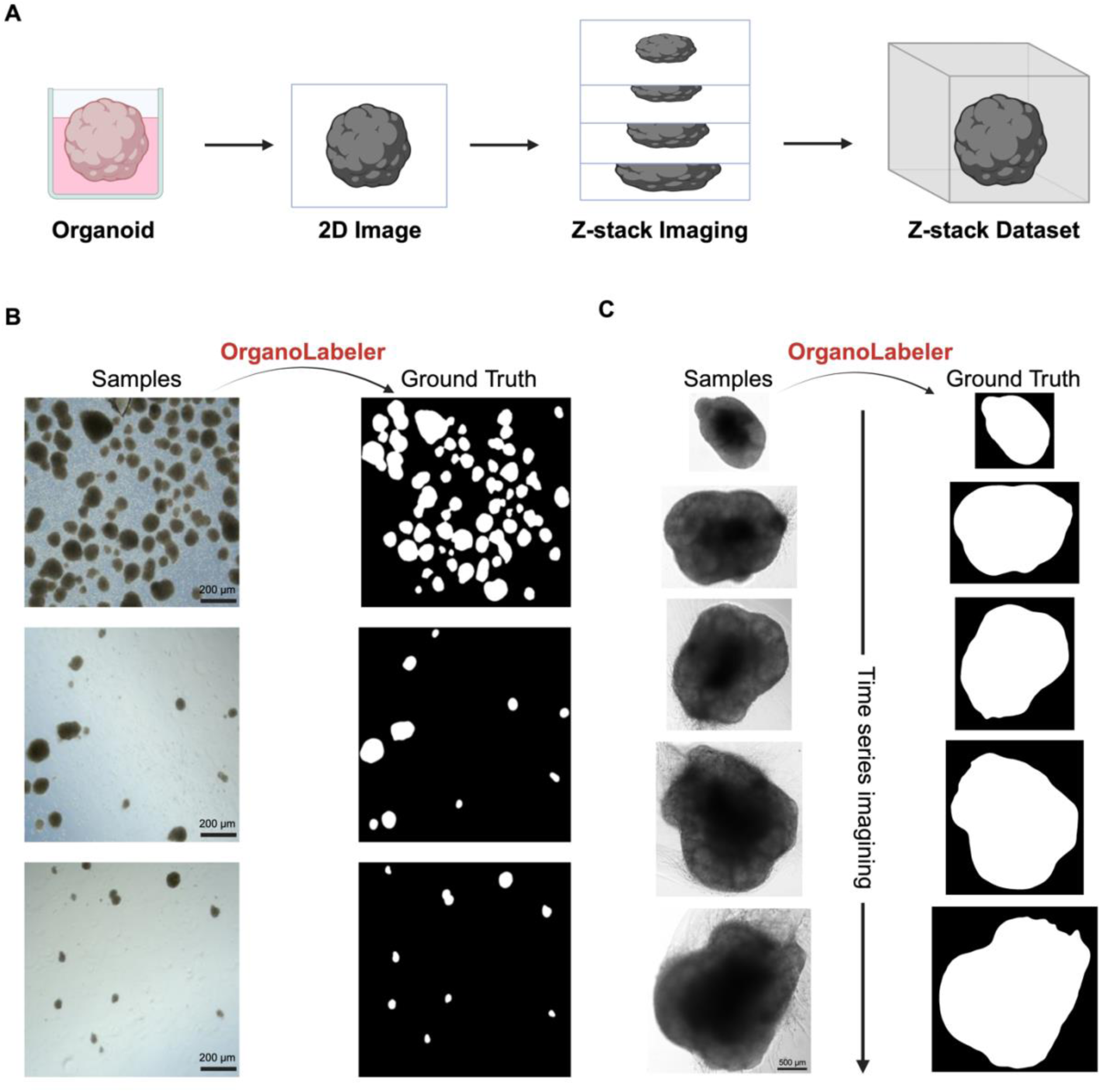
Dataset generation workflow for BrAIn. A) Z-stack imaging methodology used to accurately capture the three-dimensional complexity of brain organoids. B) Automated annotation of the embryoid body dataset using the OrganoLabeler tool. C) Time-series image acquisition for the developmental analysis of brain organoids, with dataset annotation performed using OrganoLabeler.

U-Net models were trained using the parameters given in Table 2. The model trained with the embryoid body dataset achieved a mean Intersection over Union (IoU) of 96.14 ± 0.022% and a mean Dice coefficient of 97.72 ± 0.002%. Initially, the U-Net model applied to the brain organoid dataset yielded an IoU of 73.17 ± 0.003% and a Dice coefficient of 77.55 ± 0.005%. To enhance model performance, a total of 898 synthetic brain organoid images were generated using the StyleGAN3 model, followed by data augmentation (Figure S4). As a result, the model trained with the augmented dataset achieved an IoU of 98.85 ± 0.008% and a Dice coefficient of 98.58 ± 0.007%. Each U-Net model was trained using a 5-fold cross-validation approach to ensure robustness and minimize overfitting.

The embryoid body dataset was generated using images captured on Day 1 and Day 4 from the same well of a 6-well plate (Figures 5A and S5C). Morphological analyses were performed by evaluating area, Feret diameter, and the number of embryoid bodies (Figure 5B). A decrease in the average embryoid body number on Day 4, along with an increase in the average area and Feret diameter, suggests that some embryoid bodies may have merged over time to form larger structures. The average area increased from 199.65 µm² on Day 1 to 572.28 µm² on Day 4 (Figure 5E). Similarly, the Feret diameter showed an increase from 18.51 µm to 31.84 µm. Following the segmentation step (Figure 5C) described above, roundness, circularity, area, Feret diameter, and perimeter measurements were obtained for the time-dependent developmental morphological analysis of brain organoids (Figure S5). Principal component analysis (PCA) based on time points and growth conditions was performed using the extracted measurement data (Figure 5D). The largest variance in the dataset was observed in PC1, represented along the x-axis. As shown in Figure S5B, the parameters that contributed most to the variance in PC1 were area, perimeter, and Feret diameter, exhibiting a negative correlation. The PCA plot further highlights the time-dependent morphological changes in brain organoid development. In the early stages (Day 20 and Day 27), brain organoids across all growth conditions were clustered on the right side of the plot, while, as maturation progressed, clustering shifted towards the left. When evaluated in terms of growth conditions, organoids cultured in the microfluidic chip system formed a distinct cluster on the positive side of PC2, primarily influenced by roundness and circularity (Figure S5).

**Figure 5.**
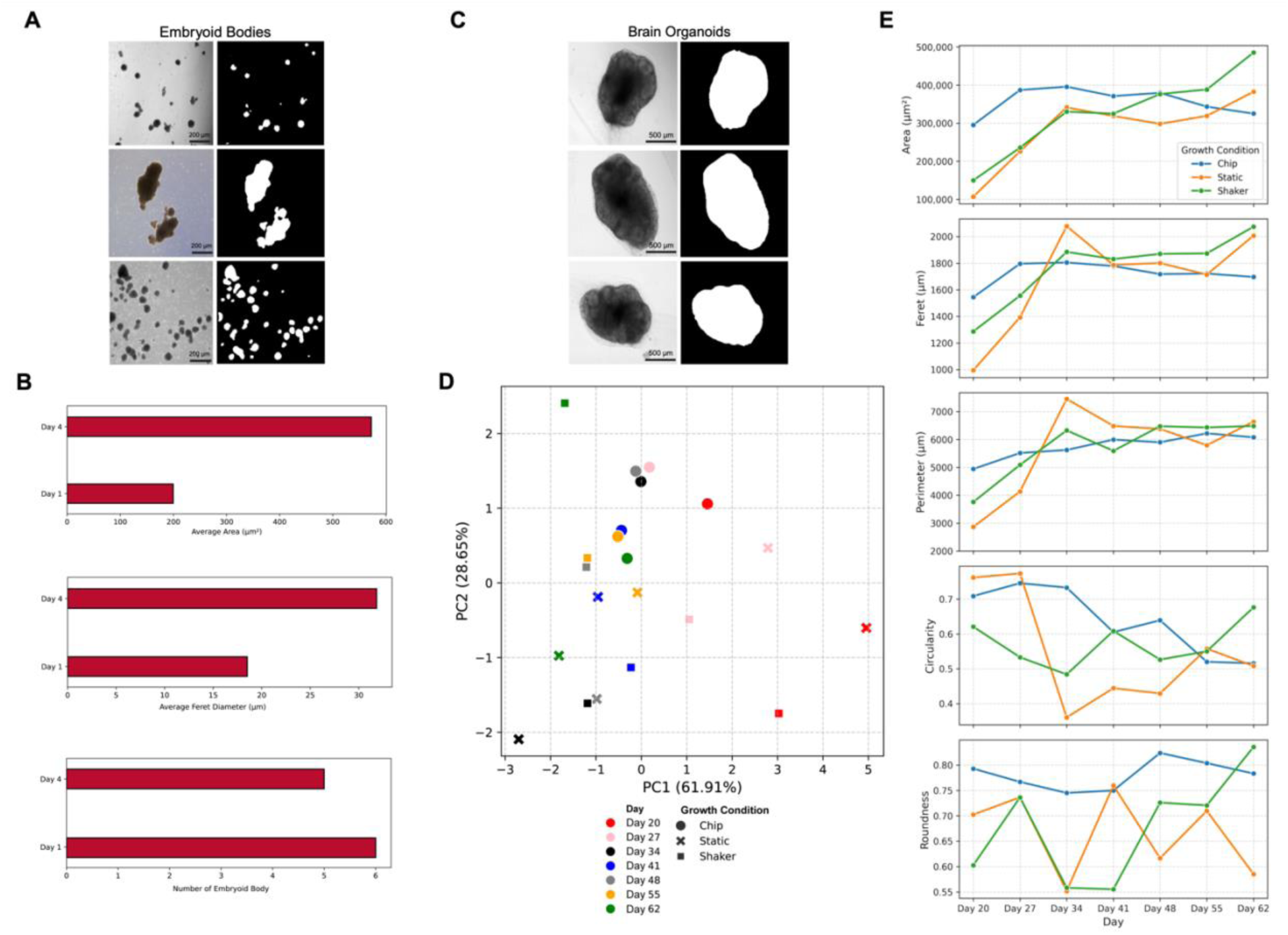
Time-series segmentation and morphological analysis of brain organoids using BrAIn. A) Representative images of embryoid bodies before and after segmentation using BrAIn. B) Quantification of embryoid body number, average area, and average Feret diameter over a 4-day period in a 6-well plate format. C) Representative raw and segmented images of brain organoids obtained from the BrAIn analysis pipeline. D) Principal Component Analysis (PCA) plot generated based on morphological parameters, including area, Feret diameter, perimeter, circularity, and roundness, to illustrate variance across different growth conditions and time points. E) Line graphs depicting the temporal progression of area, Feret diameter, perimeter, circularity, and roundness values of brain organoids under static, orbital shaker, and microfluidic chip-based growth conditions.

The time-dependent development of brain organoids under different growth environments was analyzed in terms of area changes using BrAIn (Figure 5E). The results indicate that the orbital shaker condition consistently promoted positive area growth dynamics. In contrast, in the microfluidic chip-based growth condition, initial positive growth stabilized around Day 34, after which an aerial decline was observed. Although growth progressed positively under static conditions, it remained lower compared to the orbital shaker condition.

The Feret diameter was also evaluated to assess the time-dependent development dynamics of brain organoids under different growth conditions (Figure 5E). The orbital shaker condition was the only environment that exhibited regular growth with minimal heterogeneity. While organoids cultured under static conditions occasionally reached values comparable to those in the orbital shaker condition, they failed to exhibit homogeneous growth patterns. On the other hand, organoids grown in the microfluidic chip system, despite achieving comparable maximum diameters, did not show sustained positive development over time. During the early stages (Days 20–34), Feret diameter values were higher under microfluidic chip system and orbital shaker conditions but lower in static conditions. Although static condition organoids outperformed the other conditions by the end of the early phase, they could not sustain these levels in the later stages (Days 48–62).

The perimeter changes of brain organoids under different growth conditions were also analyzed using BrAIn (Figure 5E). No significant differences were observed in perimeter measurements across all three growth conditions throughout the developmental process. Furthermore, no major effects of growth conditions were detected in the early and late developmental stages. Brain organoid development was assessed by analyzing the effects of different growth conditions on circularity and roundness values (Figure 5E). Circularity analysis revealed that brain organoids exhibited a more circular morphology across all growth conditions during the early stage. In the maturation phase, morphological heterogeneity increased gradually in the microfluidic chip system, whereas a sharper increase was observed in the orbital shaker and static conditions. In terms of roundness, brain organoids that initially exhibited a more heterogeneous morphology under orbital shaker conditions transitioned to a more homogeneous morphology in the microfluidic chip and orbital shaker systems as they matured, while increased heterogeneity was observed in static conditions.

BrAIn was further validated using the brain organoid dataset generated by Schröter et al. [74]. The segmentation performed with BrAIn achieved a Dice score of 84%, demonstrating its reliability. This dataset included brain organoids derived from four different cell lines—one healthy control and three patient-derived lines. These results indicate that BrAIn successfully analyzed an independent brain organoid dataset with acceptable accuracy, and the morphological analysis results were consistent with the published findings [74]. Time-dependent morphological analysis (Days 2, 16, and 30) of three randomly selected organoids from each group was performed using BrAIn (Figure 6A, Figure S6).

**Figure 6.**
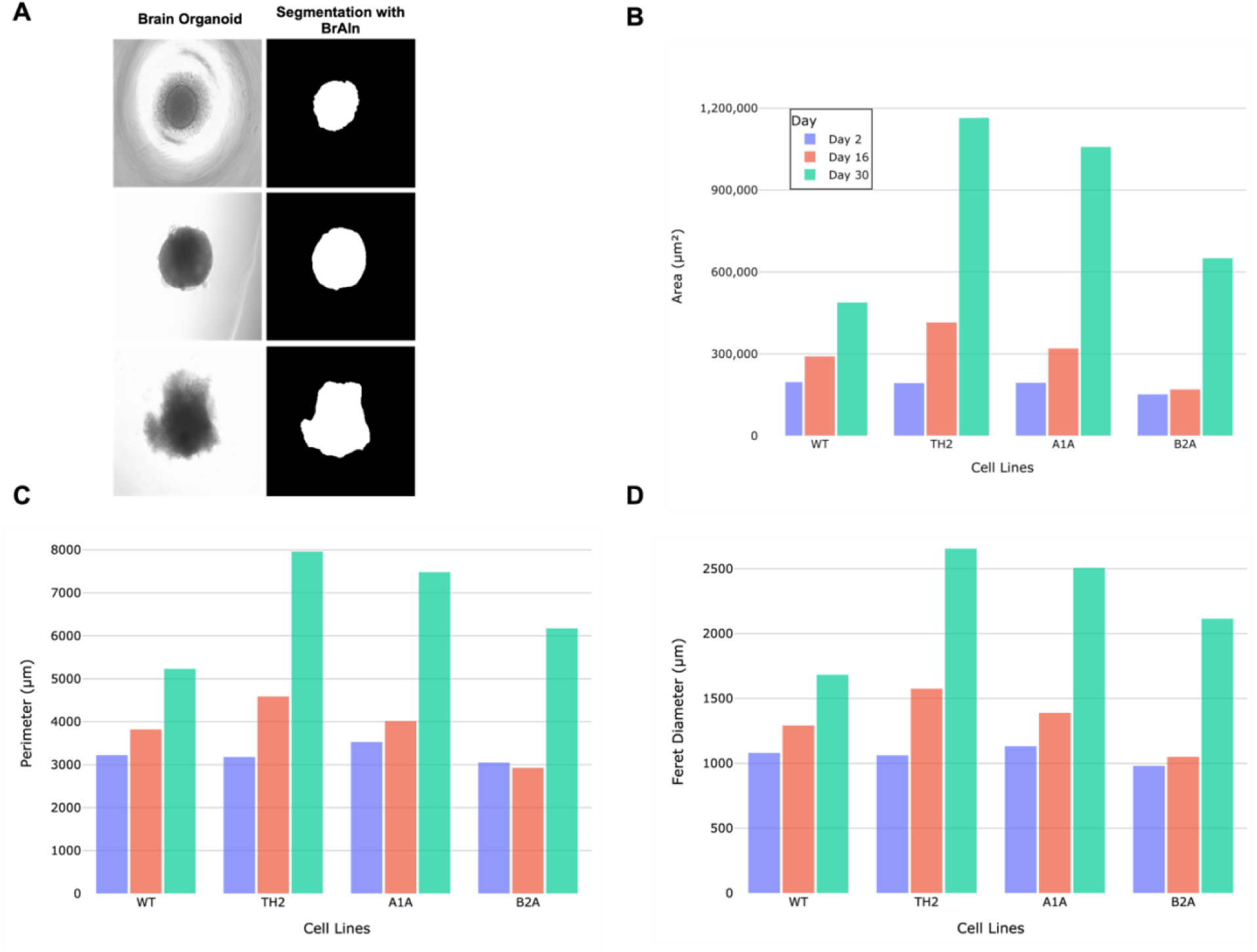
Segmentation and measurements of brain organoids generated using four different cell lines (WT, TH2, A1A, and B2A) by BrAIn. A) Representative raw and segmented images of brain organoids generated from the WT cell line (healthy control). B) Time-dependent changes in organoid area across different cell lines. C) Cell line-specific time-dependent perimeter changes of brain organoids. D) Time-dependent variations in Feret diameter across different cell lines.

Upon examining area changes, although the healthy control cell line exhibited a consistent increase over time, it lagged behind organoids derived from patient cell lines by Day 30. All patient-derived cell lines demonstrated a general trend of positive growth, with organoids generated from the TH2 cell line achieving the largest area (Figure 6B). A similar trend was observed for perimeter and Feret diameter, where all cell lines showed overall growth progression (Figure 6C-D). Notably, TH2 organoids displayed the highest values on Day 30, whereas B2A organoids followed a developmental trajectory similar to the healthy control cell line.

Roundness and circularity analyses revealed minimal variations among organoids generated from the four cell lines (Figure 6SB). However, organoids derived from the A1A cell line exhibited a more heterogeneous morphology compared to the others. Conversely, organoids generated from B2A and TH2 cell lines initially exhibited homogeneous morphology but showed increased heterogeneity over time. In contrast, the morphology of organoids generated from the healthy control cell line remained relatively homogeneous (Figure 6SB). Circularity analysis indicated initial heterogeneity in the B2A and A1A cell lines, which gradually attained a more homogeneous morphology, whereas the remaining cell lines maintained their homogeneous structure throughout the maturation process (Figure 6SB).

In order to evaluate the performance of BrAIn, we compared it with two existing organoid morphology analysis tools, OrganoID [45]] and OrgaExtractor [46], using our brain organoid dataset (Figure S8). OrganoID achieved a segmentation performance with a mean Intersection over Union (IoU) of 35% and a Dice coefficient of 46%, whereas OrgaExtractor yielded significantly lower scores, with an IoU of 2% and a Dice coefficient of 4%. Additionally, these tools were evaluated using our embryoid body dataset (Figure S8). In this dataset, OrganoID achieved an IoU of 44% and a Dice coefficient of 58%, while OrgaExtractor exhibited considerably lower performance, with an IoU of 5% and a Dice coefficient of 9%. These results demonstrate that BrAIn provides significantly higher segmentation accuracy compared to existing tools, further underscoring its robustness and reliability across different organoid datasets.

### 3.4. Neural rosette detection

Neural rosettes, critical structures in iPSC-derived neuron generation protocols, are challenging to detect visually in 2D culture systems. Their accurate identification is essential for ensuring the efficiency and reproducibility of differentiation processes. BrAIn is capable of detecting these structures using the YOLOv8 object detection model [46], achieving an initial performance of 67% mean Average Precision (mAP), 62% recall, and 70% precision with a training dataset consisting of 80 images. To enhance detection performance, data augmentation was applied by generating 293 synthetic images from real data using the StyleGAN3 model [64]. The augmented dataset, comprising a total of 373 images, significantly improved the model’s performance, achieving 96% mAP, 94% recall, and 98% precision (Figure 7).

**Figure 7.**
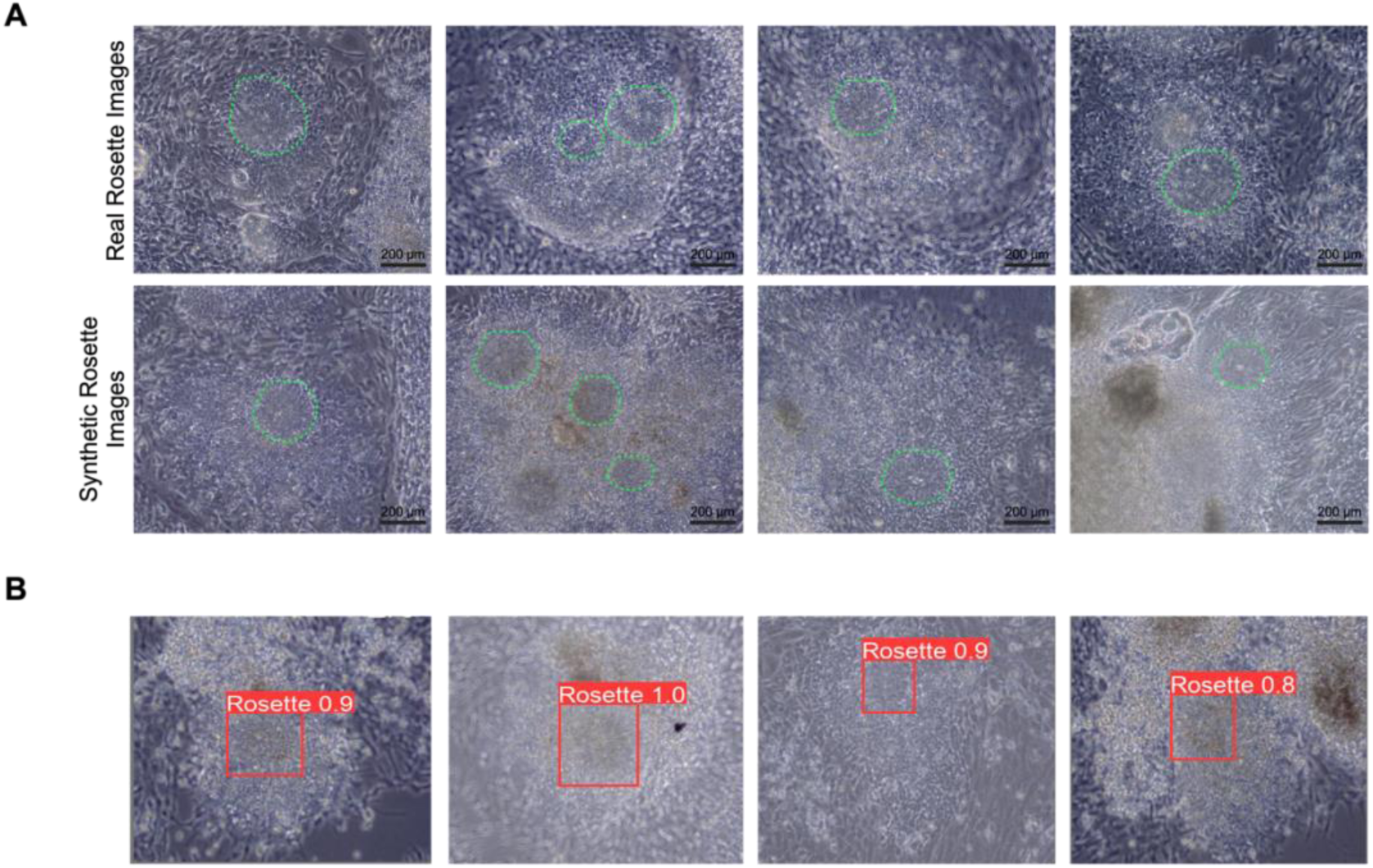
Detection of neural rosettes from iPSC-derived 2D neural culture images. A) Real neural rosette images alongside synthetic neural rosette images generated using a deep learning model. Green dashed lines indicate identified neural rosettes. B) BrAIn’s detection results for neural rosettes in 2D neural culture images.

## 4. Discussion

Morphology is one of the key indicators of a healthy and functional organoid. Understanding, monitoring, and controlling organoid morphology is among the focus of leading research groups in the field [75–78]. Morphological complexities of brain organoid typically arise from perturbations during developmental phases, resulting in more native-like brain tissue [76]. Perturbation can be achieved by carrying out development in different growth conditions [6, 8, 79] or by medium modifications [76]. The 3D nature of brain organoids enables them to be grown either under static or dynamic culture conditions [6, 63, 79]. Orbital shakers and microfluidic chip systems have positive effects on structural and functional properties in brain organoid development and better represent the human brain morphologically [7, 8]. Being able to better analyze brain organoid morphology is quite critical in terms of understanding and improving its functional and structural properties [76]. However, morphology analyses are quite complex to be done manually and time-consuming to be done semi-automatically. In this study, we developed the BrAIn tool that analyzes and quantifies the developmental process of brain organoids. BrAIn can perform three fundamental computer vision tasks: classification, segmentation, and object detection. It classifies normal, abnormal, and budding morphologies in the transition from embryoid body to brain organoid, quantifies morphological perturbations under diverse culture conditions, performs segmentation to illuminate developmental phases, and detects neural rosette structures. Although other computational tools for organoid morphology exist, BrAIn is the first to combine these tasks under three distinctive culture conditions.

The pluripotency and organoid development abilities of iPSCs in this study have been confirmed in many studies [62, 72]. Embryoid bodies are 3D cell residues that can differentiate into three different germ layers: ectoderm, endoderm, and mesoderm [72, 80, 81]. Embryoid bodies were confirmed with ectoderm, endoderm, and mesoderm markers, and their differentiation capacity was clarified (Figure S2). Organoids grown in all conditions were phenotypically confirmed for expressing brain tissue characteristics (Figure 2). Many studies have shown that brain organoids contain SOX2-expressing neural progenitors and TUJ1-expressing neurons [6, 63, 82]. We confirmed these cells in organoids developed in three different growth conditions. Neural rosettes, formed during ectodermal differentiation, express N-cadherin, indicating neural cell organization [83]. These rosettes were verified for structural integrity and neural differentiation (Figure S2).

Image-based morphological analyses have been utilized in organoid research to develop lineage differentiation protocols and assess drug responses [27, 38–56, 58–61]. Early prediction of healthy brain organoids helps avoid prolonged differentiation of abnormal samples, reducing labor and costs. Although deep learning models assist wet lab research, studies beyond organoid classification remain limited. Deep learning models are used to identify stem cells [33–36], and while morphological feature-based classification exists for some organoid types [38–42], no such classification has focused on brain organoids. BrAIn successfully predicted budding–normal and abnormal–normal categories with 92.75% and 96.83% AUC, respectively (Figure 3). It detects target features (Figure 3D) and classifies budding organoids by their budding regions, shown by Grad-CAM heatmaps. Smoother, more oval shapes are predicted as budding, while sharp boundaries indicate normal samples; in abnormal samples, BrAIn identifies irregular boundaries.

Deep learning models have been more frequently used for segmentation of organoids [43–49, 54, 55], including neural organoids [57–61]. Albanese et al. performed automatic 3D ventricle segmentation of cerebral organoids with a U-Net model (97.2% Dice) [57]. Brémond-Martin et al. used K-means and SVM to characterize early brain organoid development [60] and later adapted U-Net to achieve 88% Dice [61]. Deininger et al. segmented brain organoid MRI images (92% Dice) [59] but did not perform morphology analysis or multi-class predictions [59].

BrAIn can segment small 3D cellular structures such as embryoid bodies, organoid precursors. Embryoid bodies are on average 10 times smaller than brain organoids and range from 50 to 500 µm. BrAIn embryoid bodies were segmented with an average IoU of 96.14% and a Dice coefficient of 97.72% and analyzed in terms of average area, Feret diameter and number (Figure 5A-B). The time-dependent changes of embryoid bodies, which are cell aggregates capable of differentiating into the 3 germ layers that provide the formation of brain organoids (Figure S2), were examined with BrAIn (Figure 5A). The decrease in the number of embryoid bodies from Day 1 to Day 4, when evaluated together with the increase in mean area and ferret diameter, indicates that small cell aggregates merge to form larger ones (Figure 5B). BrAIn segmented brain organoid images collected at 7-day intervals between day 20 and day 62 from organoids cultured under three different growth conditions, including the microfluidic chip system, orbital shaker, and static culture, starting from day 16. We evaluated the growth conditions in terms of size, geometry, and surface complexity using BrAIn (Figure S5). BrAIn performs brain organoid segmentation with a 98.85% mean IoU and a 98.58% mean Dice coefficient. Our tool evaluates the effect of growth conditions on time-dependent organoid development with area, Feret diameter, perimeter, roundness, and circularity parameters. Data augmentation for the segmentation part was performed by creating synthetic images with a generative AI model, StyleGAN3 (Figure S4). To reflect the three-dimensional structure of brain organoids to the maximum extent, images were captured with the z-stack method.

By segmenting images (Figure 5C) and measuring area, Feret diameter, perimeter, roundness, and circularity (Figure 5), BrAIn reveals how dynamic 3D culture conditions—microfluidic chip and orbital shaker—compared to static culture. Dynamic 3D culture stimulates mechanotransduction, providing a more native-like microenvironment [8, 9]. We have custom designed microfluidic system for brain organoids, supplying continuous shear stress through administrated laminar media flow (Figure S1). Compared to turbulent flow generated within orbital shaker and static conditions, microfluidic chip system environment resulted in distinct clusters of morphological parameters of brain organoids at all developmental time points of PCA plot (Figure 5D). While roundness and circularity contributed positively to the largest variance of the PCA plot, size measurements contributed negatively (Figure S5). PCA1 represents the variance of organoids according to their physical size, and we found that dimensional measurements created a certain variance in different growth conditions (Figure S5).

Accordingly, organoids in wells of similar size exhibited parallel growth trends, while those in microfluidic chips showed limited area changes (Figure 5E). Organoids under orbital shaking reached the largest area, possibly due to turbulent flow facilitating nutrient exchange [6]. Feret diameter and perimeter changes paralleled area differences. The microfluidic chip system displayed more homogeneous, stable growth; circularity and roundness varied more in static and orbital shaker cultures (Figure 5E). Brain organoids grown under orbital shaker and static conditions were maintained in 9.6 cm^2^ (a well of six well plate), whereas the microfluidic chip system offered ∼1.1 cm^2^. We previously demonstrated that organoid growth area affects gene expression [9]. Accordingly, organoids in wells of similar size exhibited parallel growth trends, while those in microfluidic chips showed limited area changes (Figure 5E). Brain organoids under orbital shaking reached the largest area, possibly due to turbulent flow facilitating nutrient exchange [6]. Feret diameter and perimeter changes paralleled area differences. The microfluidic chip system displayed more homogeneous, stable growth; circularity and roundness varied more in static and orbital shaker cultures (Figure 5E).

To increase generalizability, BrAIn was also evaluated with a different brain organoid dataset (Figure 6). The LAB A subset of the dataset [74] created by Schröter et al. was segmented by BrAIn, achieving an 84% Dice score, comparable to or better than MOrgAna and OrganoSeg. BrAIn’s area-based measurements closely matched the growth performance reported by Schröter et al., highlighting its applicability to different brain organoid datasets. We also compared BrAIn with the organoid morphology tools OrganoID and OrgaExtractor (Figure S8). OrganoID offers a user-friendly interface but requires running a code block, while OrgaExtractor lacks an interface entirely. In contrast, BrAIn does not require code and is more accessible. On our brain organoid dataset, OrganoID had 35% mean IoU and 46% Dice, while OrgaExtractor had 2% IoU and 4% Dice. In the embryoid body dataset, OrganoID reached 44% IoU and 58% Dice, and OrgaExtractor 5% IoU and 9% Dice. BrAIn scored 98.85% IoU and 98.58% Dice with brain organoids, and 96.14% IoU and 97.72% Dice with embryoid bodies. Although those tools claim general applicability, BrAIn outperformed them on our datasets.

BrAIn provides comprehensive analysis for brain organoids and 2D neuron cultures, but a few limitations remain. Shadows near well-plate borders can lower segmentation performance in embryoid body images, though dense cell residues did not affect accuracy. Additionally, the number of replicates for morphology analyses at different magnifications is limited, partly due to difficulties in capturing repeated images from complex microfluidic systems.

Published tools focusing on organoid morphology analysis [43–52, 54–58, 60, 61] can handle disease modeling and drug effects but are often not user-friendly for non-coding users. By contrast, BrAIn has a straightforward interface (Figure S9). Many tools focus on a single task, whereas BrAIn covers classification, segmentation, and object detection, employing widely used models in each domain. With this study, we also present five datasets (Table 2). These brain organoid images—collected across different magnifications and times using z-stack methods— encompass early developmental stages and initial maturation. Compared against the literature, BrAIn has shown acceptable success in analyzing developmental processes of brain organoids.

The BrAIn results may lead to many future studies. In addition to the morphological effects of growth media, omics analyses can be integrated to provide a broader view of how culture conditions influence organoid size and morphology. Different growth media can be evaluated to determine the most optimal conditions for organoid development. Various microfluidic chip designs and flow rates have different effects on organoid development and yield [7, 8]; morphological analyses could guide chip design refinements. Since different orbital shaker speeds may affect the development of organoids [6, 76, 84, 85], images obtained with BrAIn could help determine the optimum shaker speed.

## 5. Conclusion

BrAIn is a powerful deep learning-based tool for the automated and objective analysis of brain organoid images. By generating quantitative morphological data and enabling direct comparisons across diverse experimental conditions, it significantly advances brain organoid research and accelerates discoveries in neuroscience and medicine. Moreover, BrAIn’s user-friendly design— requiring no coding expertise—facilitates the comprehensive evaluation of organoid development under various growth conditions, making it a versatile and invaluable asset for studying organoid biology.

## 6. Acknowledgements

This work is supported by Dokuz Eylül University ADEP TSA 2023-3026 project. B.K. is fellow of TÜBİTAK 2211C and TÜBİTAK 2250 scholarship program. E.P. is fellow of YÖK 100/2000, TÜBİTAK 2211A, and 2250 scholarship programs. A.E.E is fellow of TÜBİTAK 2210A scholarship program. The synthetic images used in this study were generated by the MeluXina Supercomputer (EHPC-BEN-2023B03-002) provided by EuroHPC JU (European High-Performance Computing Joint Undertaking).

## 8. Author Contributions

B.K and E.P contributed equally to this work. B.K., E.P. and S.G. established the hypothesis and study design. B.K. and E.P. generated brain organoids, embryoid bodies and neural rosettes. B.K and A.E.E created data sets. B.K. created BrAIn and carried out the computational experiments. B.K. and Y.B. trained AI models. B.K. and H.G trained GAN model and generated synthetic images. All authors contributed the manuscript writing. S.G., Y.B. and G.K edited the manuscript. All the authors read and approved the manuscript.

## 9. Datasets

- Embryoid Body Dataset: https://www.kaggle.com/datasets/burakkahveci/embryoid-body-dataset
- Brain Organoid Segmentation Dataset: https://www.kaggle.com/datasets/burakkahveci/brain-organoids-segmentation-dataset
- Brain Organoid Classification Dataset 1: https://www.kaggle.com/datasets/burakkahveci/brain-organoid-normal-and-abnormal-dataset
- Brain Organoid Classification Dataset 2: https://www.kaggle.com/datasets/burakkahveci/brain-organoid-budding-dataset
- Neural Rosette Dataset: https://www.kaggle.com/datasets/burakkahveci/neural-rosette-dataset
- BrAIn – setup: https://drive.google.com/drive/folders/1A9zp2cJTVHnqOcY5AAIJzMTc-oqk365f?usp=share_link

