## Supplemental Information for "BrAIn: A comprehensive artificial intelligence-based morphology analysis system for brain organoids and neuroscience"

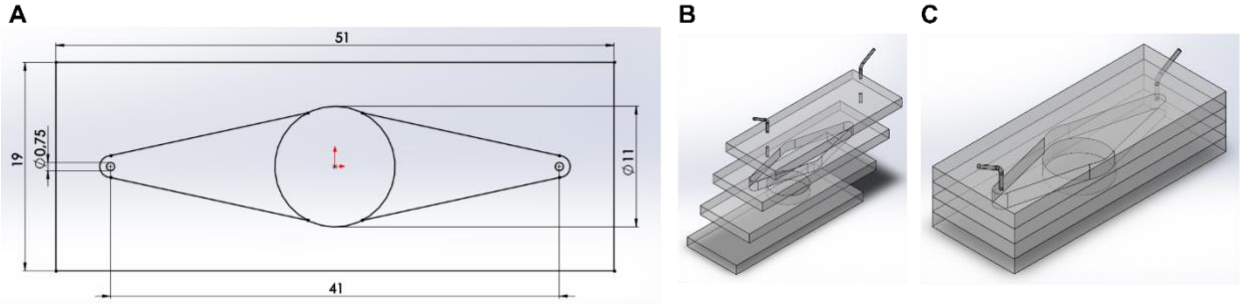


**Supplementary Figure 1.** 3D structure of the chip. A) 2D Schematic Explanation. The dimensions of the chip are specified as follows: the long side is 51 mm, the short side is 19 mm, the inlet and outlet diameters are 0.75 mm, the distance between them is 41 mm, and the organoid chamber diameter is 11 mm. B) 3D Schematic Explanation. The layers of the chip are shown as illustrated. The layers are made of PMMA, each with a thickness of 3 mm. The chip consists of a total of 4 layers. Double-sided adhesive tape was used to assemble the layers. The inlet and outlet holes are located on the top layer, and the tubing is symbolically placed in the illustration. C) 3D Schematic Explanation. The layers of the chip system are represented in the schematic drawing as shown.

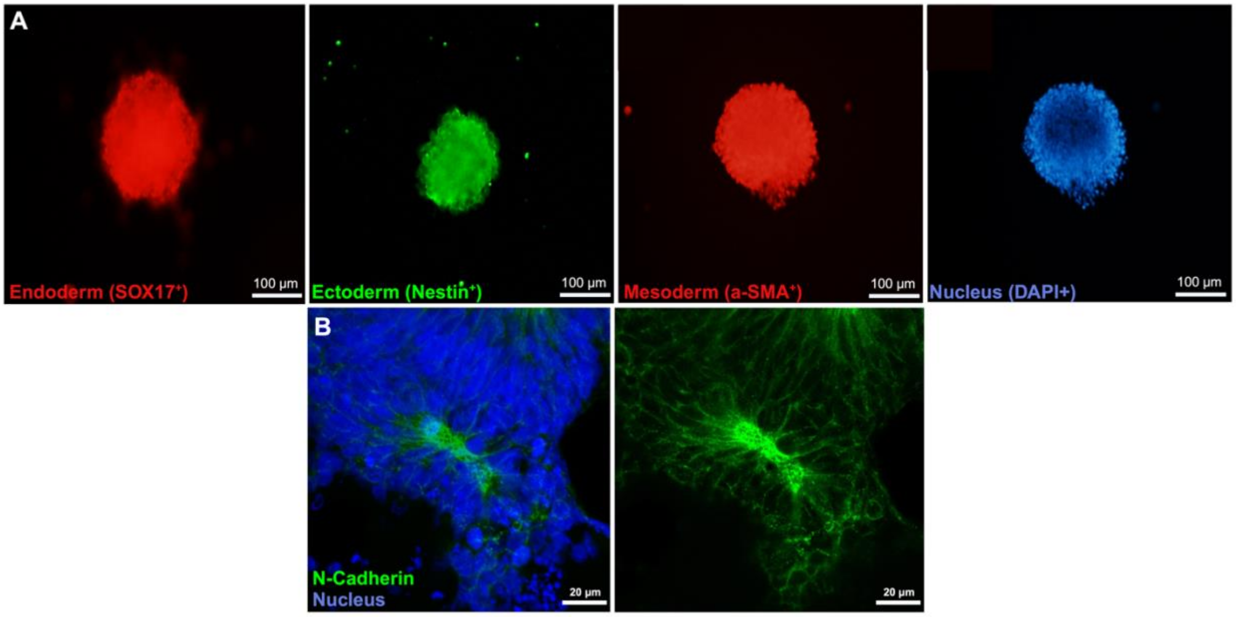


**Supplementary Figure 2.** Immunofluorescent characterization of embryoid bodies and neural rosettes. A) Embryoid bodies were stained to show the presence of three germ layers: endoderm, ectoderm and mesoderm. B) Neural rosettes were stained with N-Cadherin.

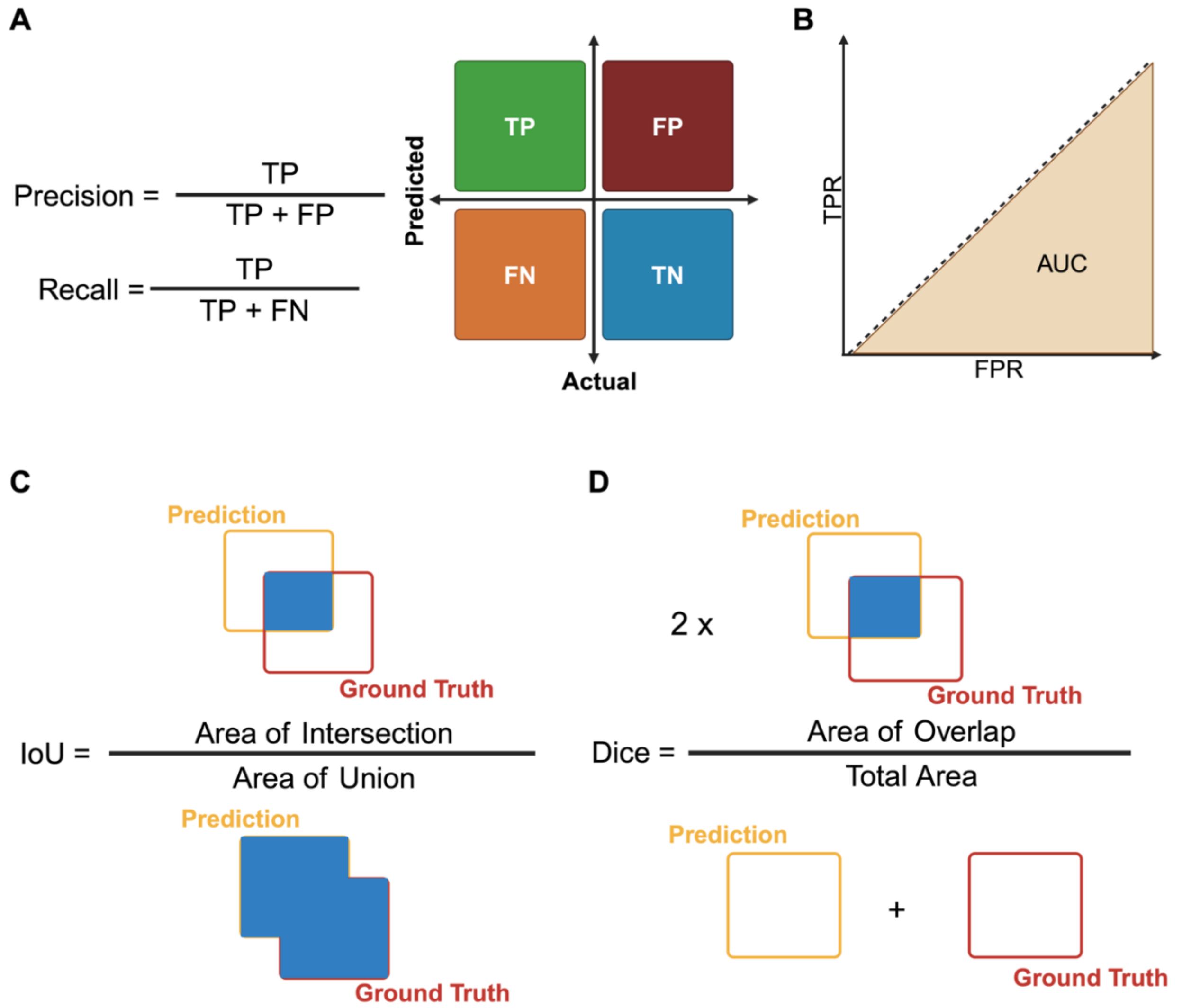


**Supplementary Figure 3.** Evaluation metrics used to evaluate BrAIn's performance. A) Mathematical calculations of precision and recall. Precision is a metric that measures the accuracy of the model in classifying an example as positive and is calculated by taking the ratio of the number of positive examples to the total number of examples classified as positive. Recall measures the proportion of true positive predictions among all true positive examples in the dataset, calculated as the ratio of true positives to the sum of true positives and false negatives. B) Definition of Area Under the Curve. Area Under the Curve is a metric that measures the performance of a binary classification model and represents the area under the ROC curve. C) Mathematical calculation of Intersection over Union. Intersection over Union is an evaluation metric used for segmentation and object detection tasks and measures the similarity between the actual segmentation mask and the segmentation output of the model. D) Mathematical definition of Dice coefficient. Dice coefficient is a measure of the similarity between two datasets and is equal to the size of the intersection divided by the sum of the sizes of the two sets

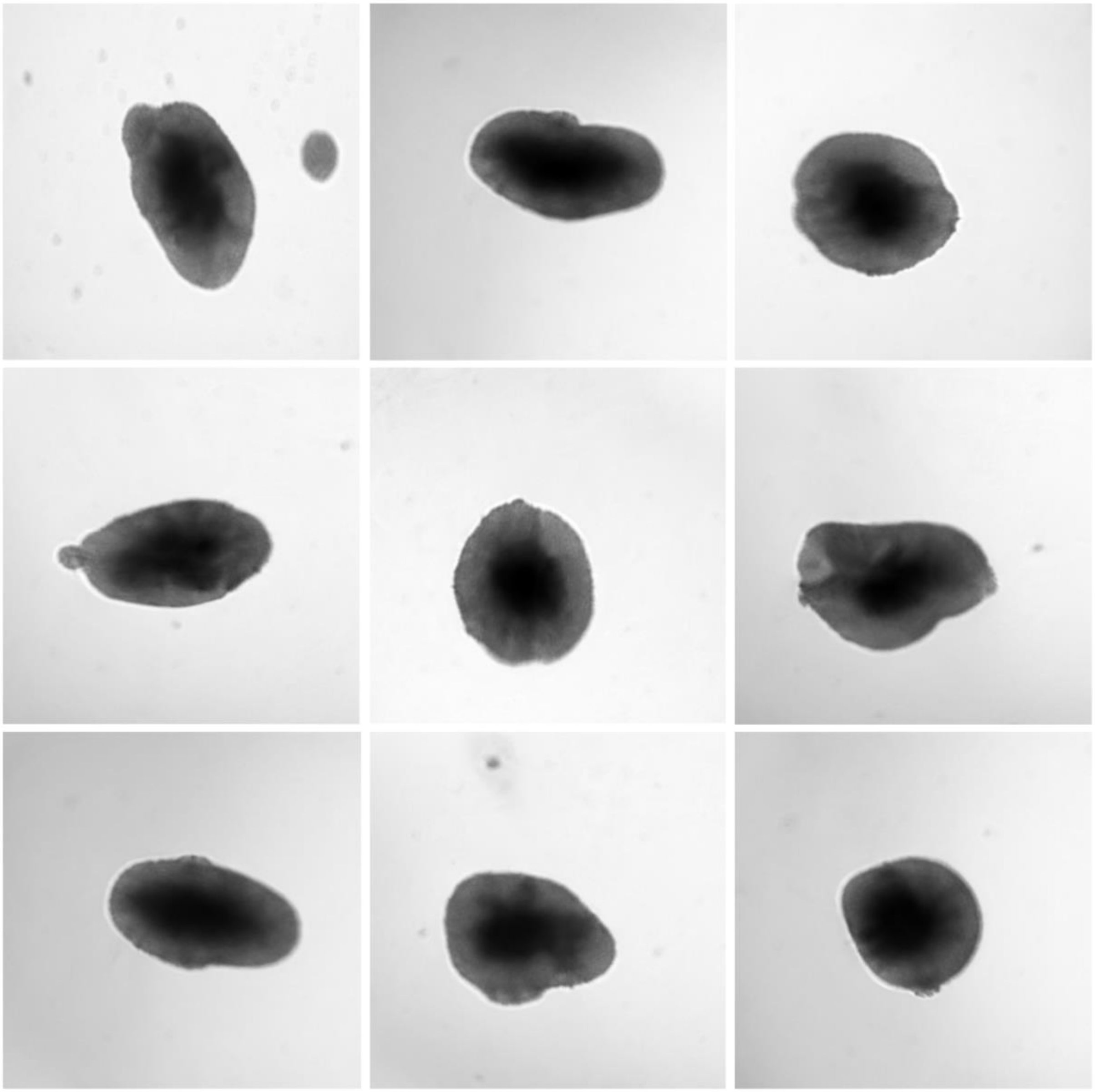

**Supplementary Figure 4.** Synthetic brain organoid images generated from real laboratory images with the StyleGAN3 model.

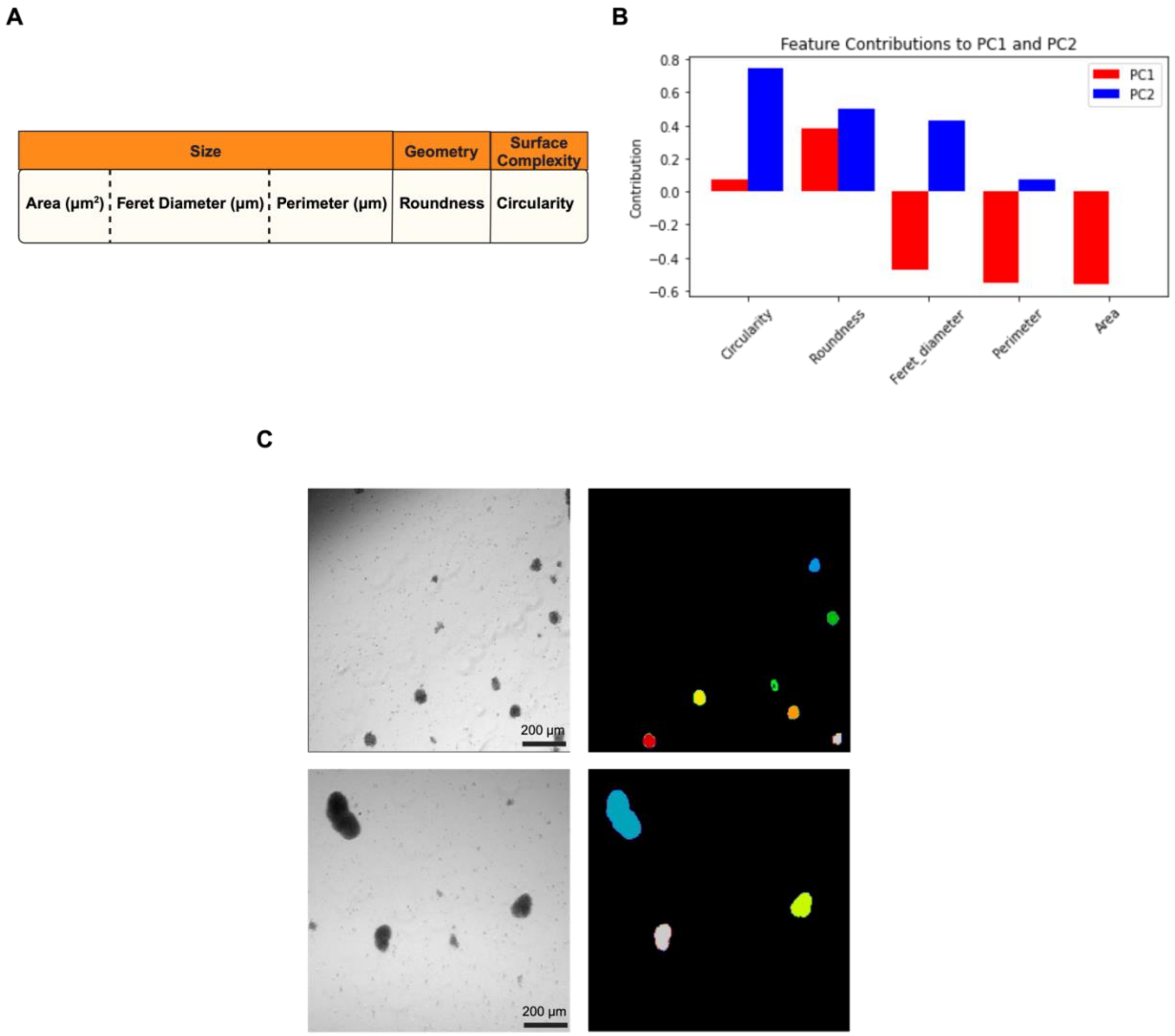


**Supplementary Figure 5.** Metrics used for morphology analysis and embryoid body segmentation. A) List of metrics used for brain organoid morphology analysis. Area refers to the total surface area of the organoids and reflects the volumetric expansion. Feret diameter measures the maximum distance between two points on the organoid and provides information about shape asymmetry. The effect of different growth conditions on spatial organization can be investigated with this metric. Perimeter corresponds to the total length of the outer border of the organoid. Roundness is a measure of how close an object is to a perfect circle geometrically. A value closer to 1 indicates more roundness. It is based on the relationship between the width and length of the organoid. Circularity describes how mathematically close an object is to a circle. It indicates the organoid's surface smoothness and overall morphological stability. B) Contribution of features to PCA plot formation. C) Embryoid body segmentation examples.

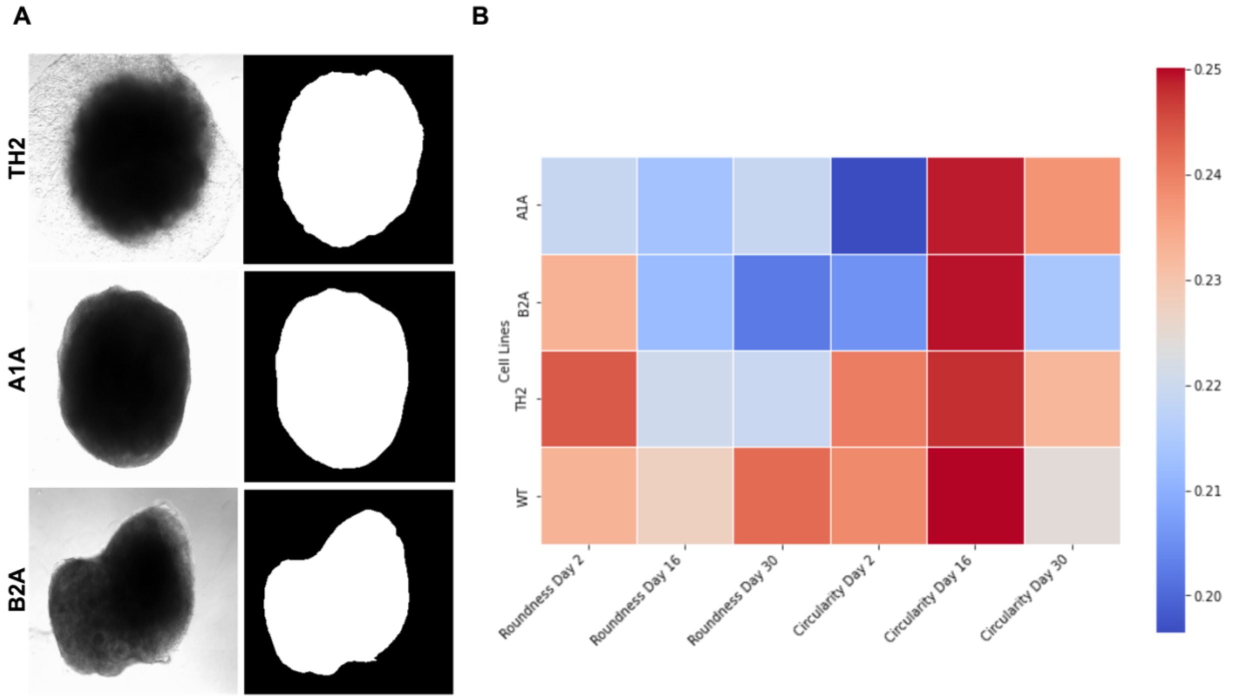


**Supplementary Figure 6.** Segmentation and morphological analysis of brain organoids generated from 4 different cell lines. A) Segmentation of brain organoids generated from three different cell lines. B) Time-series roundness and circularity analysis of brain organoids.

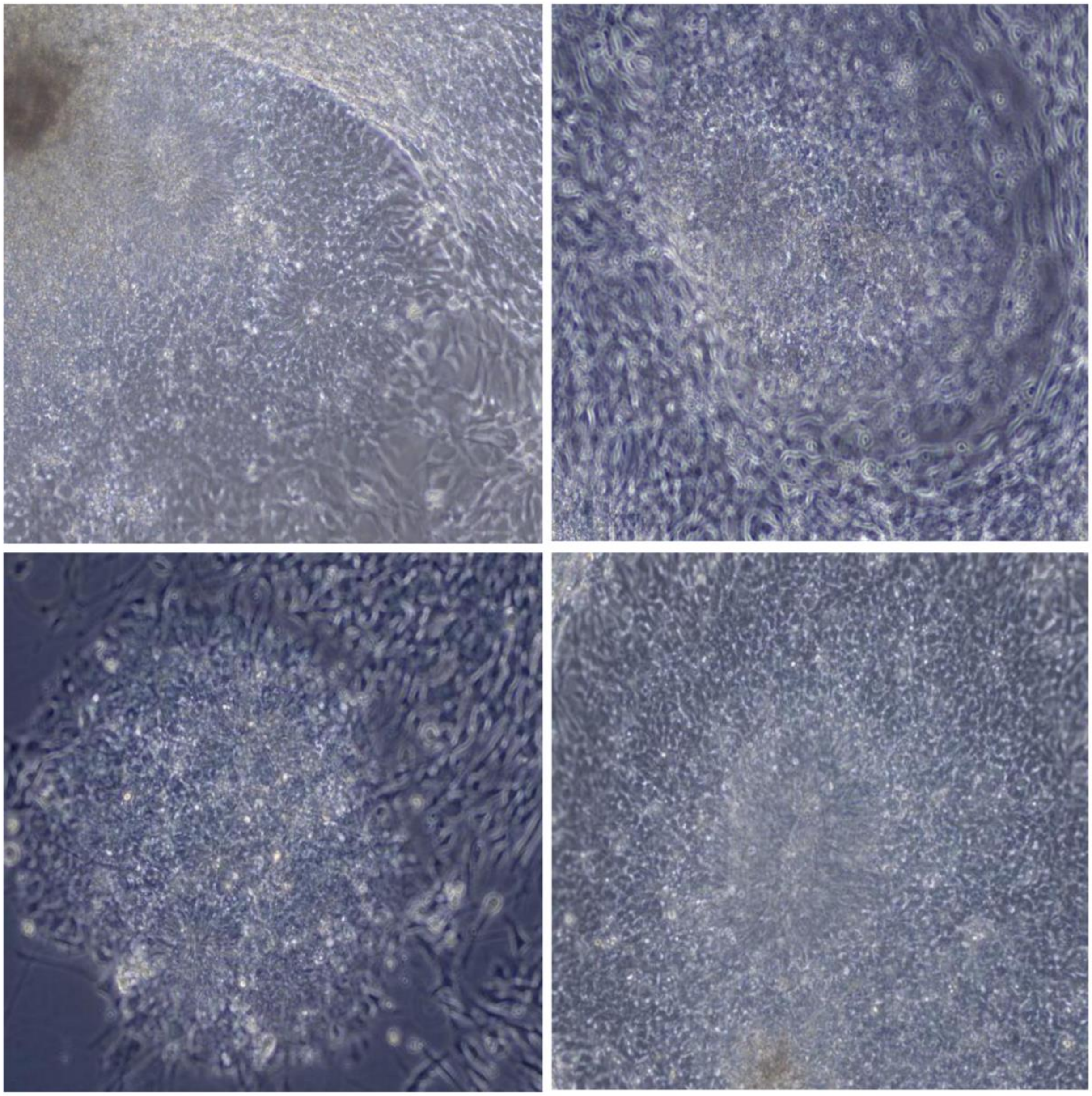

**Supplementary Figure 7.** Synthetic neural rosette images generated from real laboratory images with the StyleGAN3 model.

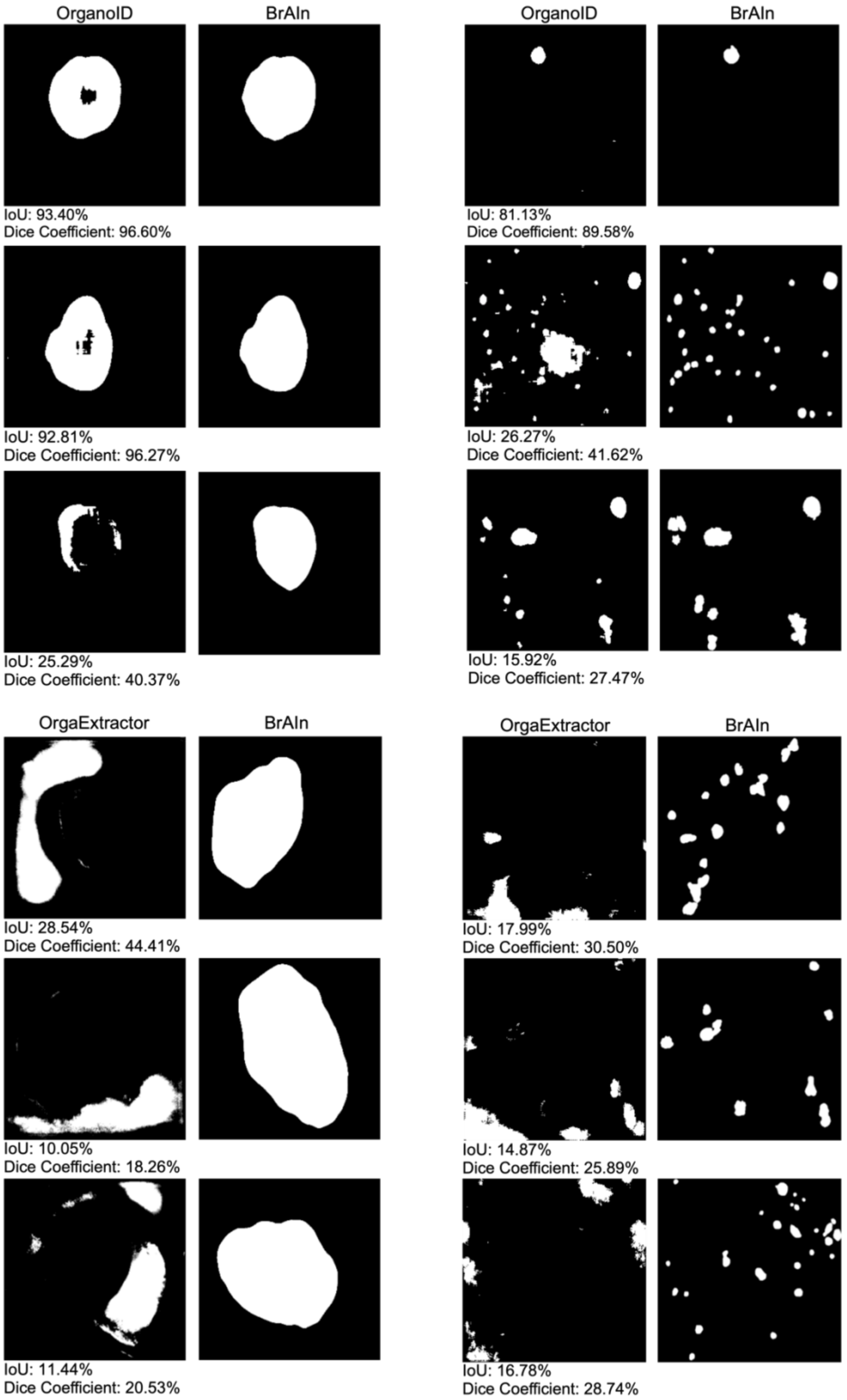


**Supplementary Figure 8.** Comparison of segmented images obtained from OrganoID and OrgoExtractor with those obtained from BrAIn

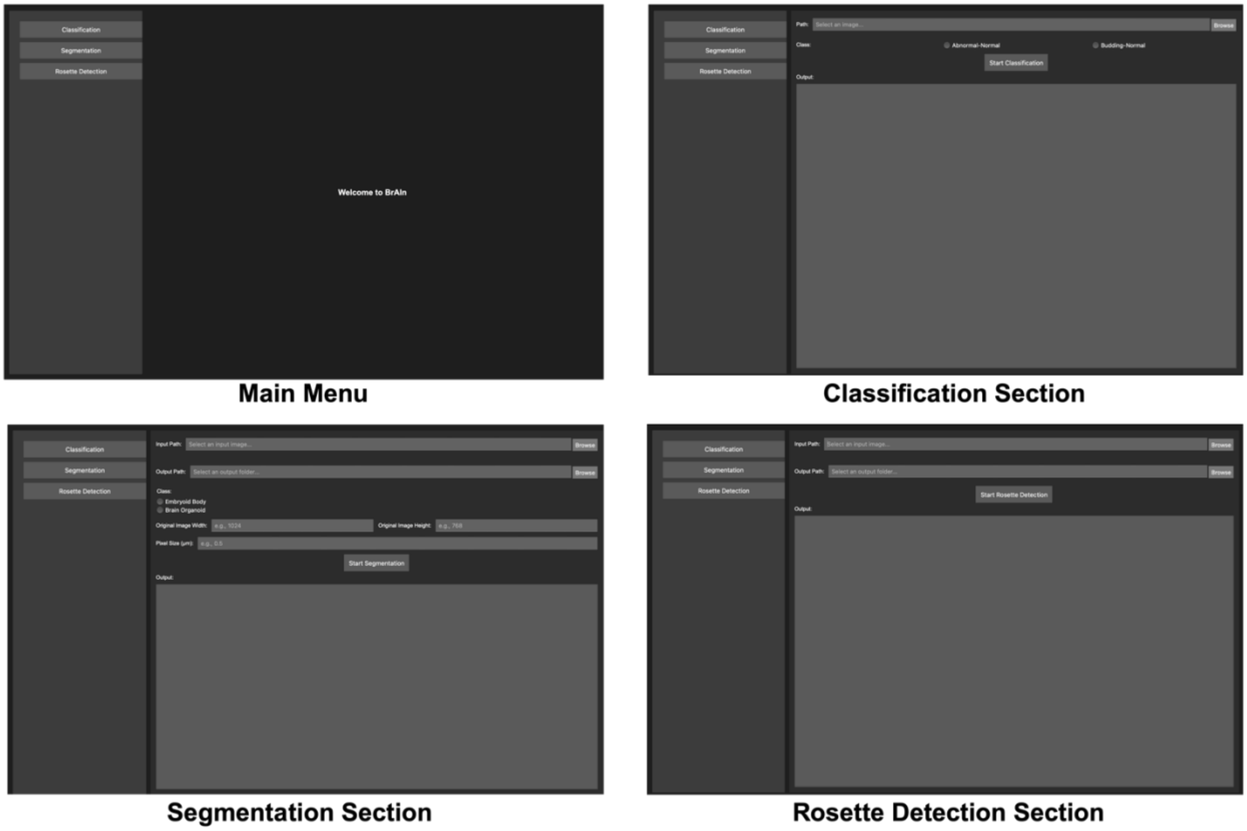


**Supplementary Figure 9.** BrAIn User Interface. The main menu contains three sections: classification, segmentation, and rosette detection. In the classification section, there are sections where the path of the image will be entered, and the classification type will be selected. When you press start classification, the result will be seen in the output section. In the segmentation section, there are input and output path sections where the path of the image will be entered, and the outputs will be recorded. Then there is the class section where the classification type is selected. There are original with, and height sections required for the morphological calculations of the image. To convert the calculated parameters of the image to micrometers, there is a Pixel size section where the number of μm in a pixel in the image taken with the relevant microscope will be entered. In the Rosette detection section, there is an input path where the path of the image will be entered and an output path where the results will be recorded. Rosette detection result information will be seen in the output section.
